# Sotrovimab retains activity against SARS-CoV-2 Omicron variant BQ.1.1 in a non-human primate model

**DOI:** 10.1101/2023.02.15.528538

**Authors:** Cécile Hérate, Romain Marlin, Franck Touret, Nathalie Dereuddre-Bosquet, Flora Donati, Francis Relouzat, Laura Junges, Mathilde Galhaut, Océane Dehan, Quentin Sconosciutti, Antoine Nougairède, Xavier de Lamballerie, Sylvie van der Werf, Roger Le Grand

## Abstract

The SARS-CoV2 Omicron variants have acquired new Spike mutations leading to escape from the most of the currently available monoclonal antibody treatments reducing the options for patients suffering from severe Covid-19. Recently, both *in vitro* and *in vivo* data have suggested that Sotrovimab could retain partial activity against recent omicron sub-lineage such as BA.5 variants, including BQ.1.1. Here we report full efficacy of Sotrovimab against BQ.1.1 viral replication as measure by RT-qPCR in a non-human primate challenge model.

## Results

Circulating SARS-CoV-2 Omicron variants have acquired mutations in the receptor-binding domain (RBD) resulting in higher ACE2-binding affinity. These changes are also associated with increasing transmission efficiency and escape to pre-existing neutralizing antibodies. In Europe, the major circulating variant BQ.1.1, derived from BA.5, is escaping most of the available anti-SARS-CoV-2 monoclonal antibody treatments. Recent *in vitro* data suggest that Sotrovimab binds Omicron subvariants, promotes Fc dependent effector functions, and still has the capacity to neutralize them (1). Moreover, treatment of S309 (Sotrovimab parent antibody; 10 or 30 mg/kg) in mice or with Sotrovimab (7 or 14 mg/kg) in hamsters provided protection from a BQ.1.1 challenge (1, 2). However, further data are needed to confirm Sotrovimab *in vivo* activity and determine whether it should remain, in addition to Nirmatrelvir-Ritonavir in the list of available treatments for patients at risk (3).

Here we report Sotrovimab efficacy in a non-human primate (NHP) model of SARS-CoV-2 infection. Three female cynomolgus macaques (*Macaca fascicularis*) aged 14-15 years and weighing between 4.6 and 7.3 kg were treated with 10 mg/kg of Sotrovimab (Xevudy) intravenously 96 h prior to viral challenge. Treated animals and two additional controls were challenged with 1×10^5^ pfu of SARS-CoV-2 BQ.1.1 (hCoV-19/France/IDF-IPP50823/2022 - EPI_ISL_15195982) via combined intranasal and intratracheal routes using an experimental protocol previously reported for other variants (4).

The treatment was carried out without any adverse effects being recorded. Sotrovimab was measured in the serum of NHP at days 1, 4, 8 and 11 post–treatment, showing a similar exposure profile (Fig 1A) as observed in humans during the COMET-ICE trial (5). Lymphopenia was reported at 2 days post-challenge (d.p.c) in non-treated animals and in 2 out of 3 treated animals as expected in this model (4). Efficacy was monitored by genomic viral RNA (gRNA) quantification using RT-qPCR (Fig. 1B). In untreated animals, gRNA was detected in tracheal fluids collected with swabs, with peak viral loads of 5.94 log10 and 5.67 log_10_ copies/mL at 2-3 d.p.c. Viral gRNA was also detected at 3 d.p.c in the broncho-alveolar lavages (BAL) at 5.25 log_10_ and 4.62 log_10_ copies/mL. By contrast, the three treated animals had viral gRNA below the limit of detection in trachea (Fig. 1B) and BAL (Fig. 1C).

**Figure 1:**
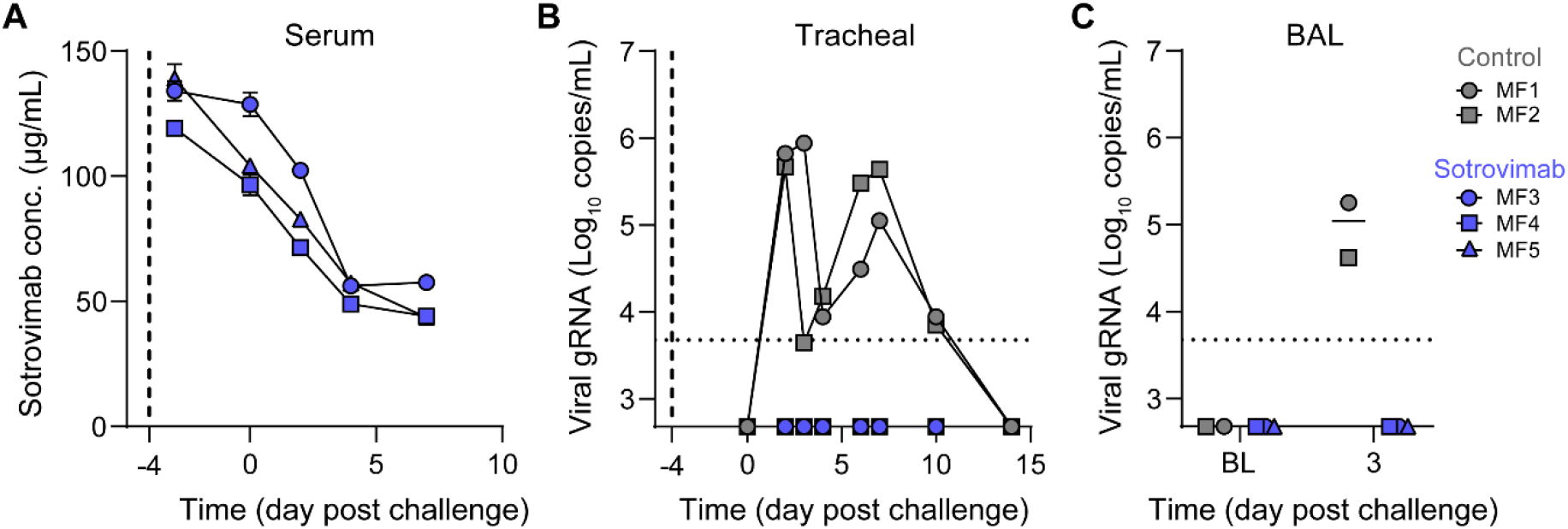
Pharmacokinetics of Sotrovimab in serum and viral loads in the respiratory tract of BQ.1.1 SARS-CoV-2 exposed cynomolgus macaques treated with Sotrovimab. Animals MF3, MF4 and MF5 were treated 4 days before challenge (blue) while MF1 and MF2 were not treated (grey). **A**. Pharmacokinetics of Sotrovimab in serum. NHPs were challenged 4 days after Sotrovimab injection. **B**. Genomic viral RNA (gRNA) was measured in tracheal fluids collected with swabs during the acute phase of infection. **C**. BAL gRNA was analyzed at baseline and day 3 post challenge. Horizontal dotted lines represent the limit of quantification (2.7 log_10_ copies/mL). Vertical dotted lines indicate the day of Sotrovimab treatment.

Sotrovimab had previously been withdrawn from the therapeutic panel due to its initially proposed poor *in vitro* efficacy against Omicron variants, in particular BA.2 (6). Here, we demonstrate that Sotrovimab completely protects NHPs from BQ.1.1 upper and lower respiratory infection. Our results thus support the use of Sotrovimab in humans against BQ.1.1, in case of ineligibility to Nirmatrelvir-Ritonavir.

## Acknowledgment

The Infectious Disease Models and Innovative Therapies (IDMIT) research infrastructure is supported by the “Programme Investissements d’ Avenir” managed by ANR under reference ANR-11-INBS-0008. The NHP experiment was part of BIOVAR and PRI programs funded by the ANRS-MIE project EMERGEN (ANRS0151)

## Authors contribution

Conceptualization: CH, RM, XDL, SVDW, RLG; Data curation: LJ, QS, NDB; Formal Analysis: CH, FT; Funding acquisition: SVDW, AN, XDL, SVDW, RLG; Investigation: FD, LJ, MG, OD, QS; Methodology: CH, RM, FD, NDB, FT, AN; Project administration: SVDW; Resources: FR; Supervision: NDB, FR, SVDW, AN, SVDW, RLG; Validation: NDB, FR, AN, XDL, SVDW; Writing-original draft: CH, RM; Writing-review & editing: CH, RM, FT, NDB, MG, FR, XDL, SVDW, RLG CH and RM are co-first authors, CH contributed to design of the study, contributed to animal work, the coordination of the experiments, data analysis and the writing of the paper.

## Appendix

## Materials and Methods

### Ethics and biosafety statement

Five female cynomolgus macaques (*M. fascicularis*), aged 14-15 years and originating from Mauritian AAALAC-certified breeding centres were included to this study. All animals were housed in IDMIT facilities (CEA, Fontenay-aux-roses), under BSL-3 containment (Animal facility authorization #D92-032-02, Préfecture des Hauts de Seine, France) and in compliance with European Directive 2010/63/EU, the French regulations and the Standards for Human Care and Use of Laboratory Animals, of the Office for Laboratory Animal Welfare (OLAW, assurance number #A5826-01, US). The protocols were approved by the institutional ethical committee “Comité d’ Ethique en Expérimentation Animale du Commissariat à l’ Energie Atomique et aux Energies Alternatives” (CEtEA #44) under statement number A20-066 and A22-006. The study was authorized by the “Research, Innovation and Education Ministry” under registration number respectively APAFIS#29191-2021011811505374 v1 and APAFIS# 36939-2022042217237124 v1

## Antibody treatment

Sotrovimab (Xevudy 500 mg solution to dilute for perfusion) is a commercial antibody also named VIR-7831 and GSK4182136. The treatment was administrated 96 h prior challenge by intravenous perfusion in 30 minutes. Sotrovimab was diluted at 1.434 mg/mL in NaCl 0.9 % and injected at a dose of 10 mg/kg, which represents a volume between 34 and 50 mL.

Animals were monitored for heart rate, respiratory rate and oximetry every ten minutes from treatment initiation until 30 min after the end of injection.

### Quantification of Sotrovimab monoclonal antibody

Sotrovimab exposure was measured using the commercial enzyme-linked immunosorbent assay (ELISA) anti-SARS-CoV-2 Quantivac (IgG) kit (Euroimmun) which is directed against the S1 domain of the spike protein.

Results were expressed in binding antibody units per mL (BAU/mL) following manufacturer instructions and converted to μg/mL using blank plasma from untreated/infected animals spiked with known quantities of Sotrovimab.

### SARS-CoV-2 challenge

Treated and control animals were challenged with SARS-CoV-2 BQ.1.1 (hCoV-19/France/IDF-IPP50823/2022 - EPI_ISL_15195982), provided by the National Reference Center for respiratory viruses at Institut Pasteur, via the combination of intranasal (1/10) and intratracheal (9/10) routes (day 0), using atropine (0.04 mg/kg) for pre-medication and ketamine (5 mg/kg) with medetomidine (0.05 mg/kg) for anesthesia. Tracheal fluids and blood samples were regularly collected following challenge. Broncho alveolar lavages (BAL) were performed at 3 and 10 days post challenge. NHPs were followed for behavior assessment and clinical score during the 7 days of the infection. Blood cell counts, hemoglobin and hematocrit were determined from EDTA blood using a DXH800 analyzer (Beckman Coulter).

### Viral quantification

Viral genomic RNA (gRNA) was quantified in swabs and BAL samples by RT-qPCR with a plasmid standard concentration range. The protocols describing the procedure for the detection of SARS-CoV-2 is available on the WHO website (https://www.who.int/docs/default-source/coronaviruse/real-time-rt-pcr-assays-for-the-detection-of-sars-cov-2-institut-pasteur-paris.pdf?sfvrsn=3662fcb6_2)

